# Family Analysis with Mendelian Imputations

**DOI:** 10.1101/2020.07.02.185181

**Authors:** Augustine Kong, Stefania Benonisdottir, Alexander I. Young

**Affiliations:** Big Data Institute, Li Ka Shing Centre for Health Information and Discovery, University of Oxford, UK; Center for Economic and Social Research, University of Southern California, Los Angeles, CA, USA

## Abstract

Genotype-phenotype associations can be results of direct effects, genetic nurturing effects and population stratification confounding. Genotypes from parents and siblings of the proband can be used to statistically disentangle these effects. To maximize power, a comprehensive framework for utilizing various combinations of parents’ and siblings’ genotypes is introduced. Central to the approach is *mendelian imputation*, a method that utilizes identity by descent (IBD) information to non-linearly impute genotypes into untyped relatives using genotypes of typed individuals. Applying the method to UK Biobank probands with at least one parent or sibling genotyped, for an educational attainment (EA) polygenic score that has an *R*^2^ of 5.7% with EA, its predictive power based on direct genetic effect alone is demonstrated to be only about 1.4%. For women, the EA polygenic score has a bigger estimated direct effect on age-at-first-birth than EA itself.

## Introduction

Standard genotype-phenotype association analyses, such as those typically performed for genome-wide association studies (GWAS), involve only the phenotypes and the genotypes of the proband. However, to separate the direct genetic effects from the indirect genetic effects and other confounding factors, in addition to the proband’s genotypes, genotypes of family members such as parents and siblings are often necessary^1^. Even though much more family data can be expected in the future, either through deliberate ascertainment or as a consequence of a substantial fraction of the population being genotyped, family data are currently somewhat limited. Thus, for now and for the future, it is important to develop methods that can get the most out of the data available. Here we consider a model where the proband’s phenotype depends on the genotypes of four people ---the proband, the parents, and one sibling. When genotypes of one or more family members are unavailable, they are treated as missing-data, and imputed in an appropriate manner. This setup serves two purposes: (a) it allows different data types to be treated under one analytic framework, increasing flexibility and power, (b) statistical efficiency is increased through non-linear imputation of the missing genotypes using the observed genotypes. Genotyped sib-pairs with untyped parents, a common data type, benefit the most from this approach. Compare to standard analyses^2–5^, our method of imputing parental genotypes, which incorporates the identical-by-decent (IBD) information between sibs, adds information and allows for the estimation of sibling genetic nurturing effect. Moreover, it includes the modelling of asymmetric sib-pairs, *e.g.* siblings of different gender, and highlights the utility of the genotypes of a sibling whose phenotype is either missing or is not directly comparable with that of the proband.

This paper is organized as follows. (i) The basic model and parameters are introduced. (ii) The fifteen possible genotype data patterns, one complete plus fourteen incomplete, are presented and (iii) the various forms of imputations, linear and IBD-based non-linear, are described and illustrated by examples. (iv) Provide conditions for the estimates obtained using data with imputations to be unbiased, *i.e.* the estimates, while having different standard errors, have the same interpretations as those obtained using complete data. We call this property *estimate consistency.* (v) Illustrate the extension from handling one phenotyped sibling (sib) to two phenotyped sibs and introduce a model that incorporates phenotypic asymmetry between sibs. A note on extension to genotyped sib-ships of size bigger than two. (vi) When the data or estimates from different missing data patterns are combined, it is a form of multivariate (parameter) meta-analysis^6^. Someimtes the results are not intuitive, *e.g.* estimates often have smaller standard errors than one expects from applying univariate principles. (vii) Extension of analyses of individual variants to that of polygenic scores and discuss the consequences of assortative mating. (viii) Empirical study based on UK Biobank data. (ix) Discussion.

## Models and Setup

The proband is defined as the person with known phenotype *Y*, the response variable. We start with a single-locus model where *Y*, conditional on the genotypes of the proband and the parents, has expectation

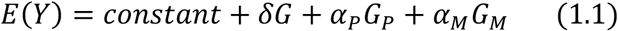

where *G* is the genotype of the proband, *G_
P_* is the genotype of the father, *G_M_* is the genotype of the mother, and *δ* is the direct effect. *Y* is treated as a quantitative variable, but it does not have to be normally distributed and indeed can be binary. As long as the variance explained by the *G*’s is small relative to the variance of *Y*, results given will apply exactly or approximately. Note that (1.1) is in effect the same as a model previous used^7^ where the explanatory variables are the transmitted and non-transmitted alleles, as the explanatory variables this model are a one-to-one linear transformation of those in the other. The parameters *α_P_* and *α_M_* can be written as

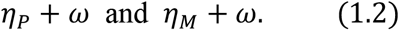

where *η_P_* and *η_M_* denote parent-of-origin (PO) specific genetic nurturing effect, and ω captures all confounding effects that have not been adjusted out, including assortative mating induced confounding. Note that, following our previous work^7^, the genetic nurturing effects of the parental alleles, *η_P_* and *η_M_*, are meant to incorporate not only the genetic nurturing effects of the parents, but also include the contributions from older ancestors and siblings. In particular, when the proband has a sibling, the model is extended to

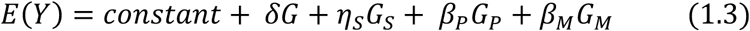

where *η_s_* denote the genetic nurturing effect of the sibling’s genotype. Because a parental allele has ½ chance of being passed onto a sibling,

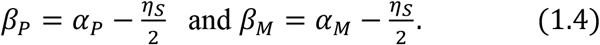

The *α*‘s are more natural parameters than the *β*’s, as the former are well defined regardless of whether the proband has any siblings. Nonetheless, the introduction of *η_s_* and the *β*’s is necessary when family analysis, in the absence of parental genotypes, is performed using the genotypes of a sibling. While (*δ,β_P_, β_M_, η_s_*) is the parameter vector being directed estimated through fitting the model (1.3), its estimate could easily be transformed into an estimate of other parameters vectors such as (*δ, α, η_M_ — η_P_, η_s_*) where

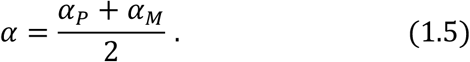

Notice that because the estimates of the four parameters are correlated, estimates of *δ* and *α* will change and usually have reduced standard errors if the assumptions that *η_P_ = η_M_* and *η_s_* = 0 are made, a question of bias-variance trade-off. Finally, it is noted that the sum (*δ* + *α*), referred to as the *population effect* here, corresponds to what is being estimated in a proband only genotype-phenotype analysis performed by most GWAS studies.

### Treatment of Missing Data

Genotypes (explanatory variables) in the complete-data model (1.3) that are unobserved are treated as missing-data. Including the complete-data case, there are 2^4^ — 1 = 15 complete-missing data patterns (Fig. 1, genotyped individuals shaded). For example, Fig. 1a, 1b, 1e, and 1h, correspond respectively to complete data, parents-proband trios, genotyped sib-pairs, and the standard GWAS *singletons* setup with proband genotyped only. Missing genotypes are imputed either linearly (○ or blank) or non-linearly (+ or ⊕). Linear imputation is predicting an unobserved genotype using a linear combination of the observed genotypes with coefficients that minimize mean squared error (MSE). With alleles coded 0 or 1 and population frequency of allele 1 denoted by *p*, the four genotypes *(G, G_s_, G_P_, G_M_*), assuming random mating, have known variance-covariance matrix (Table 1a). Even though the *G*’s are not normally distributed, the formulas for best linear predictions are the same as those established for multivariate normal variables. However, as illustrated below, based on the known non-normal joint distribution of the G’s, sometimes a missing genotype can be imputed as a non-linear function of the observed genotypes with higher correlation with the actual genotype than any linear imputation.

**Figure 1:**
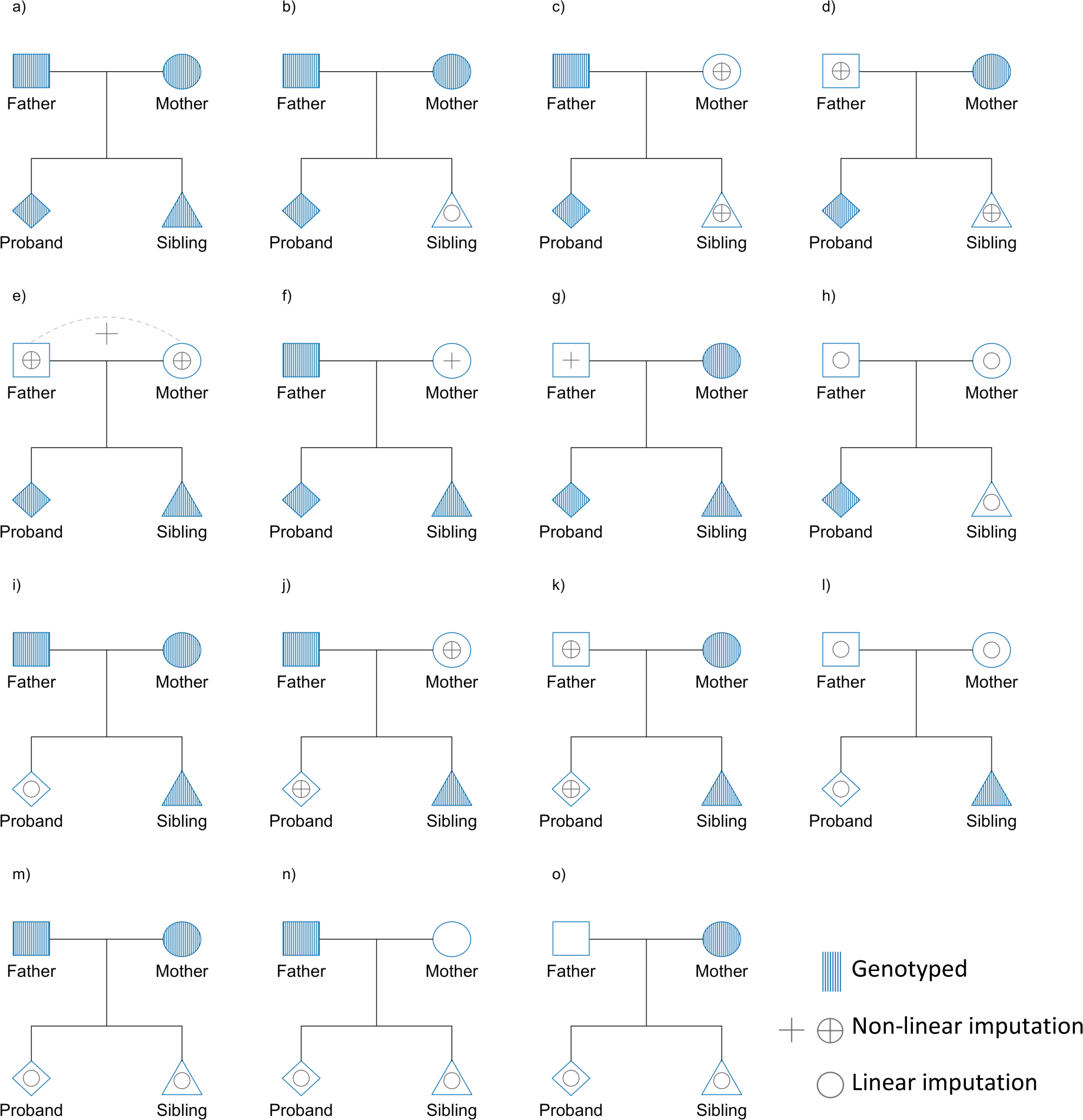
Mendelian imputation for 15 different missing data patterns for nuclear families. Proband refers to an individual with phenotype information. Shaded individuals are directly genotyped. ○ denotes linear imputation from observed genotypes, + denotes non-linear imputation from observed genotypes and IBD information, ⊕ denotes non-linear imputation in the case where the resulting covariance matrix of the four genotypes (observed and imputed) is not of full rank (see Table 1). The genotypes of the blanked mother in 1n and blanked father in 1o are imputed by a constant, the population frequency.

**Table 1.**
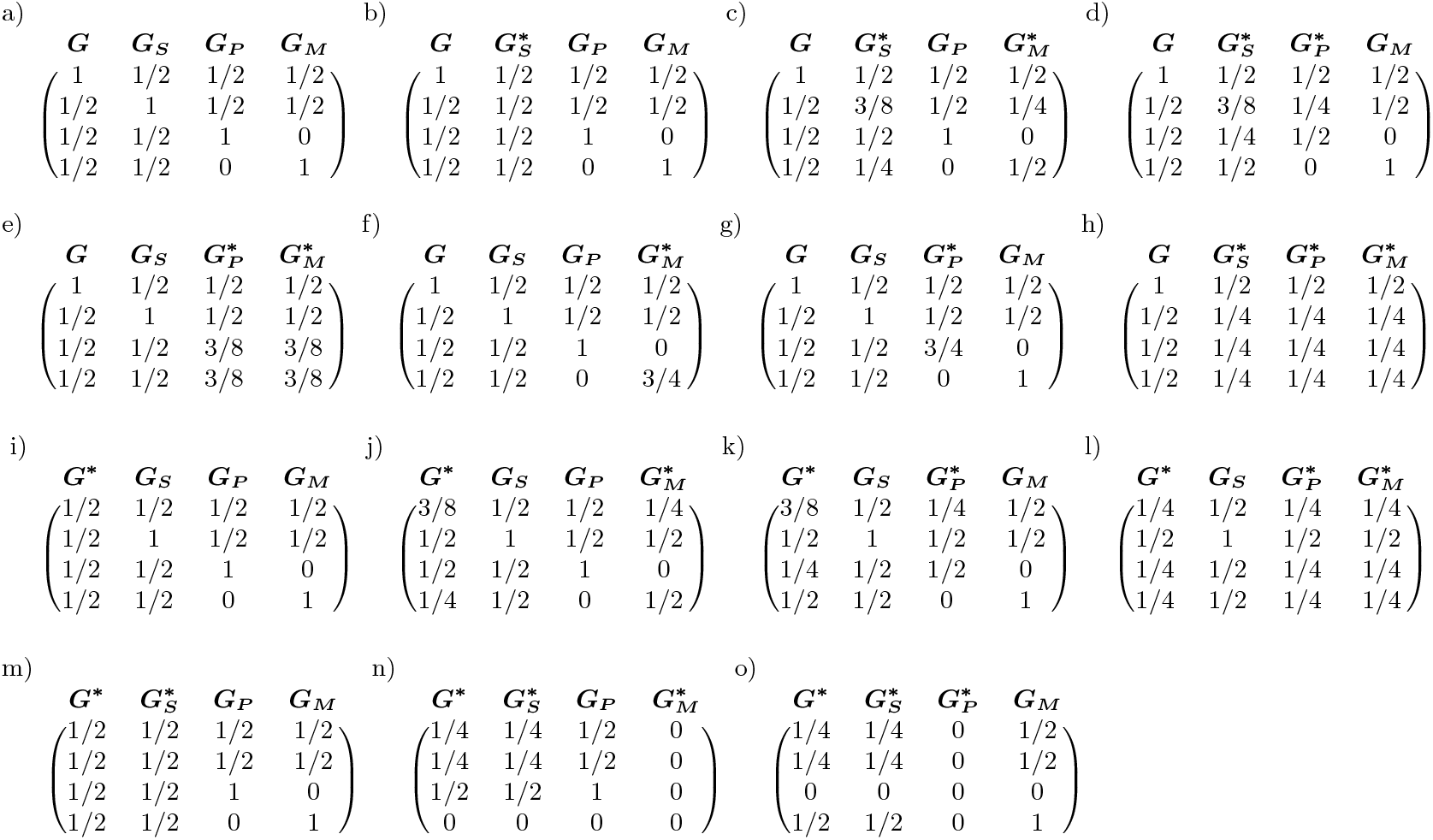
Mendelian imputation and resulting scaled variance-covariance matrices. Each matrix shows the variance-covariance structure within the nuclear family given observed and imputed genotypes. The labels a) to o) correspond to the those in Fig. 1. *G* denotes proband’s genotype, *G_s_* denotes sibling’s genotype, *G_P_* denotes father’s genotype and *G_M_* denotes mother’s genotype. * indicates an unobserved genotype that is imputed (either linearly or non-linearly) using observed genotypes. Displayed are 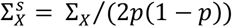. It is scaled so that the diagonal entry corresponding to an observed genotype is 1.

There are seven cases in Fig. 1 where IBD-based non-linear imputations are possible. However, with respect to the joint distribution of the genotypes, up to symmetry, there are only 3 distinct equivalent classes --- (1c,1d,1j,1k), (1e), and (1f,1g). One case from each of the equivalent class is covered below.

#### Mother and proband genotyped

In Fig. 1d, *G* and *G_M_* are observed, but not *G_P_* and *G_s_*. Based on the known variance-covariance matrix of the genotypes,

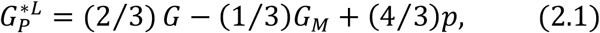

is the best linear predictor of *G_P_* (formula for computing linear imputations in Supplementary Information), with 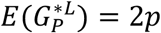, and 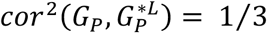. *G_P_* can be decomposed as *T_P_* and *NT_P_*, denoting respectively the allele transmitted to the proband and the allele not transmitted. If *T_P_* is known, the best and natural prediction of *G_P_* is

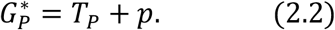

satisfying 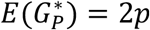, and 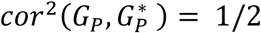. *T_P_* is equal to *G* — *T_M_*, where *T_M_* is the allele transmitted from mother to proband. Given *G* and *G_M_, T_M_* is known unless *G* and *G_M_* are both heterozygotes. In that case, unless the target SNP is very close to a recombination event in the mother-proband meiosis, *T_M_* can be deduced through a phased neighbouring SNP which is homozygous for one member of the mother-proband pair and heterozygous for the other (Fig. 2).

**Figure 2:**
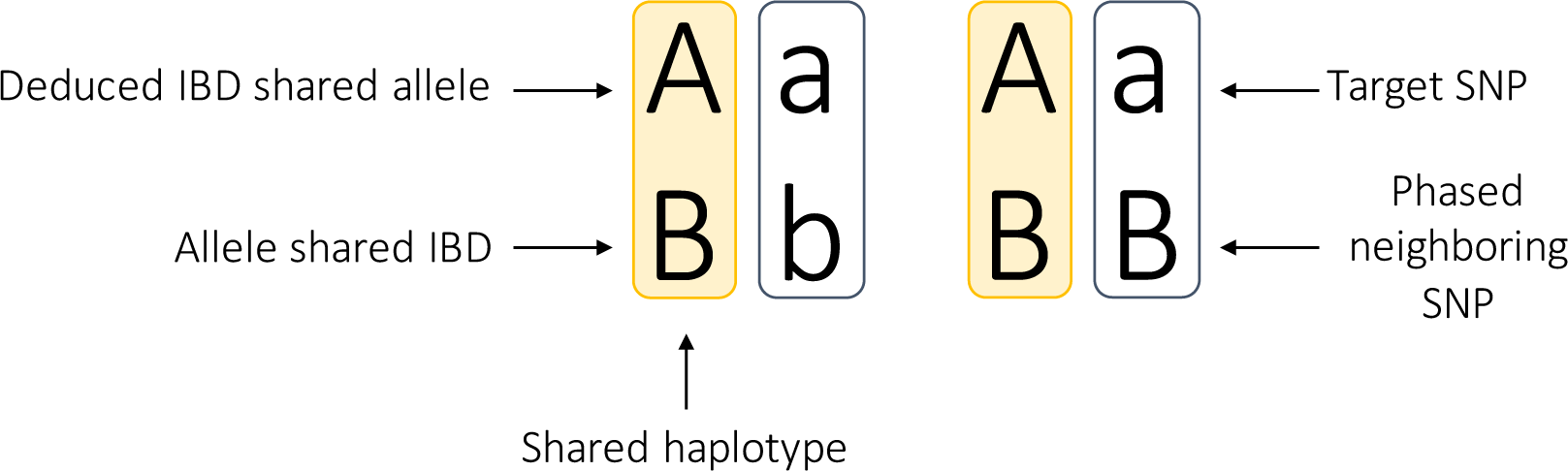
How the allele shared IBD between two individuals can be determined when both are heterozygous at the target SNP. This applies to a parent-offspring pair, who always shared one allele IBD, and a sibling pair at locations where they are determined to be sharing one allele IBD. A neighbouring SNP which has been phased with the target SNP, and is homozygous for one sib and heterozygous for the other is employed to resolve the uncertainty. For the individual on the left above, the B allele must be the allele shared with the individual on the right. Thus through the phased haplotype A-B, it is determined that allele A, as opposed to a, is the shared IBD allele.

To avoid the phasing step, with some loss of information, one could impute *G_P_* by *2p* when *G* and *G_M_* are both heterozygotes^8^, an event occurring with probability *p*(1 — *p*). Relative to linear imputation, the increased correlation between *G_P_* and 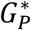 leads to increased information for parameter estimation. If data with this pattern are analysed on its own, as opposed to mixing together with other patterns, then linear imputation corresponds to no imputation at all. In particular, if *Y* is regressed on *G* and *G_M_* only, it can be shown that the fitted coefficients have expectations [*δ* + (2/3)*α_P_*] and [*α_M_* — (1/3)*α_P_*] respectively. Without the assumption that *α_M_ = α_P_*, it is not possible to obtain an unbiased estimate of *δ*. By contrast, by regressing *Y* on *G, G_M_* and 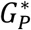, in which case the fitted coefficients would have expectations *δ, α_M_* and *α_P_* respectively, the same as if *G_P_* is observed and included in the regression. However, replacing *G_P_* by 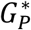 would not just change the variances of individual parameter estimates, it would have an impact on the whole variance-covariance structure of the correlated estimates. Also, if the assumption *α_M_ = α_P_* is made, then *δ* can be estimated with or without non-linear imputations, but the variance of the estimate obtained with imputation is only 3/4 of that of the estimate obtained without imputation. If data with this missing data pattern are analysed jointly with other data in a single regression with all four explanatory variables included, then 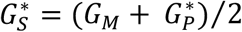. However, even though 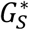 is a non-linear function of *G* and *G_M_*, it is a linear function of *G_M_* and 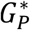. Thus, if data of this pattern are analysed by themselves, with 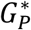 already included in the regression, adding 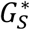 would only introduce collinearity (indicated by ⊕ in Fig. 1).

#### Siblings genotyped only

In Fig. 1e, *G* and *G_S_* are observed, but not *G_P_* and *G_M_*. Without PO information, the best linear prediction of *G_PM_ = G_P_* + *G_M_* is

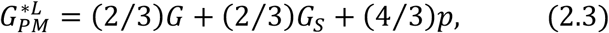

with 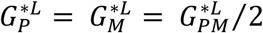. At an autosomal locus, two siblings share 0, 1, or 2, alleles IBD with probability 1/4,1/2, and 1/4 respectively. With the large number of SNPs included in any of the recent genome-wide genotyping arrays, given their genotypes, the number of alleles shared IBD between a specific sib-pair at a specific locus can usually be determined quite accurately^9^ unless the locus is close to one of the paternal or maternal recombination events for the siblings, which happens infrequently. With IBD number known (Fig. 3), *G_PM_* can be nonlinear imputed as

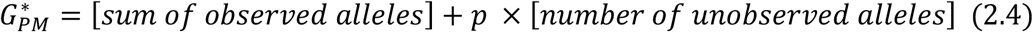

with 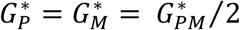. The number of unobserved parental alleles is equal to four minus the IBD number, and observed parental alleles are summed without double counting. Similar to the *parent-proband* case (Fig. 2), when IBD = 1 and both siblings are heterozygous at the target SNP, there is uncertainty as to which is the IBD shared allele. This can again be resolved through a neighbouring phased SNP which is homozygous for one sib and heterozygous for the other. If phasing is not performed, with some loss of information, *G_PM_* can be imputed as 1 + 2*p* in this situation^8^. Assuming the double-heterozygous situation is resolved through phasing, it can be shown that 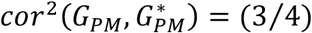, an increase over 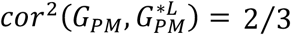. If sib-pair data are analysed by themselves, since 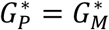, it is natural to regress *Y* on *G*, *G_S_* and 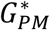. The respective fitted coefficients are then unbiased estimates of *δ,η_S_*, and *β* = (*β*_P_ + *β_M_*)/2. By contrast, if *Y* is regressed on *G* and *G_S_* only, the respective fitted coefficients have expectations [*δ* + (2/3)*β*] and [*η_S_* + (2/3)*β*] respectively. By taking the difference of these two fitted coefficients, one can obtain an unbiased estimate for (*δ — η_S_*). Without imputations, *δ* cannot be estimated without bias unless the assumption *η_S_* = 0 is made. Even then this estimate is suboptimal: when only one sib is a proband, *i.e.* phenotyped, it has a variance *4/3* times that of the estimate of *δ* obtained with imputations and conditioning on *η_S_* = 0 (see below for the double-proband case).

**Figure 3:**
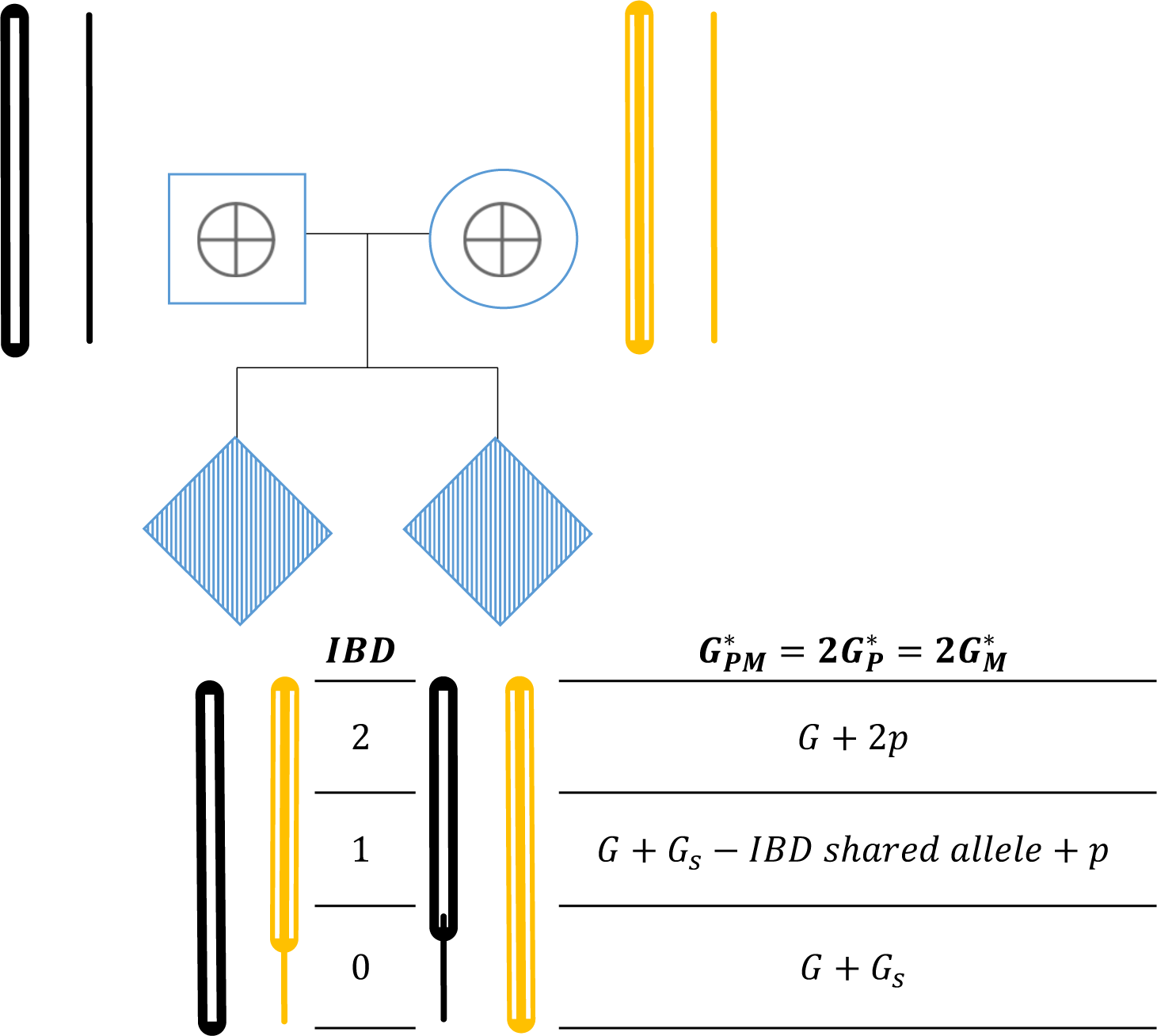
Mendelian imputation of parental alleles given the IBD status of genotyped sibling pairs. Parent-offspring pairs share one allele IBD at each locus. Siblings share 0, 1 or 2 alleles IBD with probabilities 1/4,1/2., and 1/4 respectively. Given array genotypes, the number of IBD alleles shared in realization can often be determined with little uncertainty. Illustrated is how (*G_P_* + *G_M_*) is imputed given IBD number

#### Two siblings and mother genotyped

In Fig. 1g, *G, G_S_*, and *G_M_* are observed. Similar to the previous two examples, if the paternal alleles transmitted to the proband and sibling are estimated to be non-IBD, then 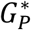 is the sum of those two alleles. Otherwise, 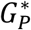 is the common paternal allele plus *p*-By regressing *Y* on *G, G_S_, G_M_* and 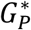, the fitted coefficients are unbiased estimates of *δ, η_S_, β_M_*, and *β_P_*, the same as with complete data.

### Estimate Consistency and Multivariate Meta-Analysis

Let 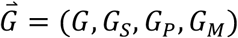 and let 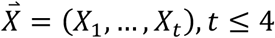, be explanatory variables used in an analysis satisfying the condition that the phenotype *Y* is correlated with 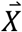 only through 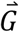, *i.e. Y* is conditional independent of 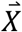 given 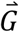. Let matrices ∑,∑_*X*_, and Σ_*cov*_, be respectively 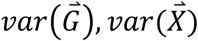, and 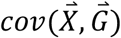. The model (1-3) can be rewritten as

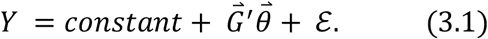

where 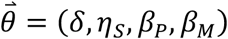 and 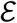 and 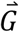 are uncorrelated. If *Y* is regressed on 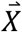 linearly, the corresponding model is

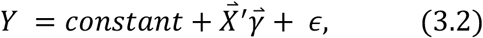

with 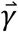 satisfying

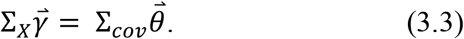

If Σ_*X*_, is of full rank, then

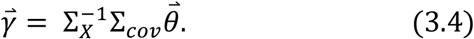

In addition to being used in earlier examples to calculate the expectations of the fitted coefficients when 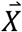 consists of a subset of the *G*’s, this formula is key to understanding the regressions performed with imputed genotypes. For the proposed imputations, Table 1 gives

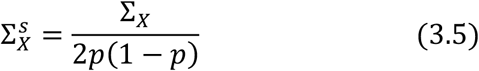

for the fifteen data patterns in Fig. 1. 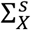, is a scaled version of Σ_*x*_, with the property that the diagonal entry of an observed genotype is one, *e.g.* instead of 0 or 1, alleles are coded as 0 or 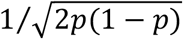. (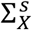 is given for linear imputations in Supplementary Table 1.) Consider the Fig. 1g example studied earlier. Here 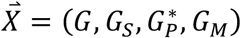. Regardless of how 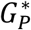 is computed, because of the overlapping variables in 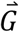 and 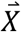, matrices Σ, Σ_*X*_, and Σ_*cov*_, are by definition the same apart from entries in row 3 and column 3, and Σ_*X*_, and Σ_*cov*_ are the same except for column 3. The imputation 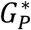 we proposed further ensures that Σ_*X*_, = Σ for all entries except entry [3, 3], where Σ_*X*_[3,3]/Σ[3,3] = 3/4 (equivalent relationships reflected by 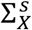, in Table 1a and Table 1g), and *Σ*, = Σ_*cov*_ for all entries (Supplementary Information). It follows that 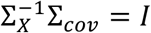 and, most importantly

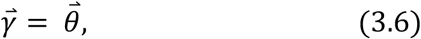

the property we call *estimate consistency*. With our proposed imputations, the relationships between the matrices extend to all fifteen patterns in Fig. 1 (Supplementary Information). In particular, entries of Σ, and Σ_*X*_, and equivalently their scaled versions, are equal except for those entries where both indexes tag imputed genotypes, and

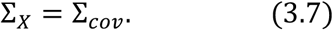

The latter also implies that for an imputed genotype, the corresponding diagonal element in 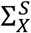 (Table 1) is the correlation-squared between actual and imputed genotype. For example, in the 1g case,

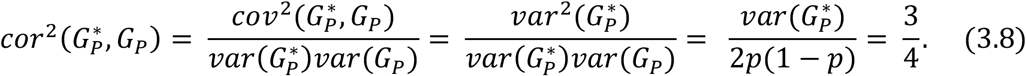

Among the fifteen patterns, seven involve non-linear imputations (+ or ⊕) and *rank*(Σ_*X*_) = number of genotyped (shaded) family members plus 1. For the others, *rank*(Σ_*X*_) = number of genotyped members. Thus Σ_*X*_ is of full rank for 1a, 1f and 1g. For the other twelve cases, if the data with any of these patterns are analysed by themselves, the number of explanatory variables would have to be reduced to eliminate collinearity, as demonstrated above for 1d and 1e. However, with data from more than one pattern, by imputing all the missing genotypes, with both linear and nonlinear imputations, a single regression can be performed with all the data mixed together. Specifically, consider data from *k* different patterns indexed by *i*. For *i* = 1, …, *k*, let *n_1_* be the sample sizes, 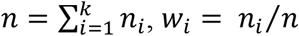, and Σ_*Xi*_ and Σ_*covi*_ be the Σ_*X*_ and Σ_*cov*_ of pattern *i*. The combined data have sample size *n* and variancecovariance matrix

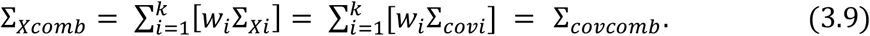

This is because Σ_*Xi*_ = Σ_*covi*_ for each *i*, and our imputations satisfy 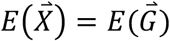 for all data patterns. If Σ_*Xcomb*_ is of full rank, then the combined data can be analysed by one regression based on the complete-data model (1.3). Notice that Σ_*Xcomb*_ would be full rank as long as we have some data from patterns 1a, 1f or 1g. It would also be of full rank if the data include pattern 1e and cases from either 1b, 1c or 1d. If Σ_*Xcomb*_ is of rank 3, and the data do not include any genotyped sibling, then model (1.1) can be considered as the complete-data model. Similarly, if Σ_*Xcomb*_ is of rank 3 and the parents’ genotypes are always missing, then fitting a model with the parental genotypes combined is appropriate. In general, individually, the different data patterns have different variance-covariance structures for the explanatory variables and through their inverses impact the variance-covariance structures of the parameter estimates.

### Both siblings are phenotyped

Here we consider the case where both siblings are phenotyped and thus both are probands. Let *Y*_1_ and *Y*_2_ denote respectively the phenotypes of sib1 and sib2, and let *G* and *G_S_* denote their respective genotypes. Let 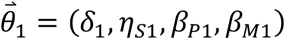, and 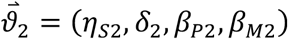. The complete genotype-data model is

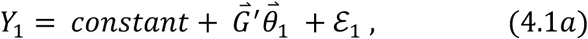

and

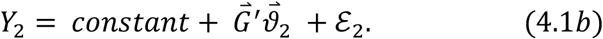

The subscript of 1 or 2 for the parameters allows for asymmetry between the sibs. For example, if sib1 is male and sib2 is female, or sib1 is the younger sib, the parameters could take on different values. Assuming *Y*_1_ and *Y*_2_ to be each standardized to have variance one, under asymmetry, 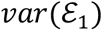 and 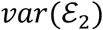 are not necessarily equal. For simplicity, we assume the difference is negligible and denote their average value as *σ*^2^. The parameters (and their corresponding estimates) can be reparametrize as averages and differences:

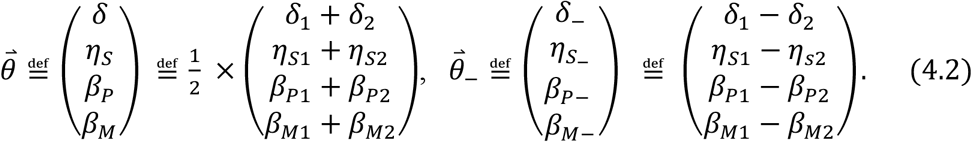

From (4.2), related parameters are similarly defined, *e.g. β* = (*β_P_ + β_M_*)/2, *α = β + η_S_*/2, and *α*_ = (*α*_1_ — *α*_2_). Assuming symmetry corresponds to conditioning on 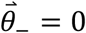. Estimates can be obtained by performing regressions that correspond to (4.1a) and (4.1b). However, because 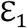 and 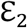 are correlated, determining the variance-covariance matrix of the parameters is more complicated. In Supplementary Information, we show how to do that by reparameterizing the responses to 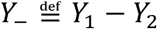 and 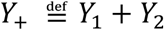. Here we focus on the case where only the siblings are genotyped, one of the most common data-type. The variance-covariance of the average parameters (Supplementary Information) are

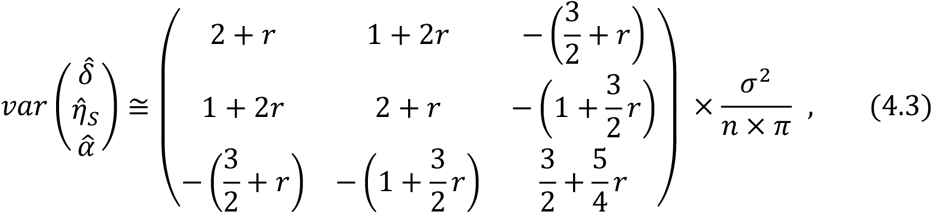

where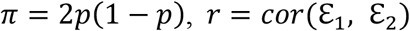, and *n* is the number of sib-pairs. The variance-covariance matrix of the estimates of the difference parameters, which are uncorrelated with the estimates of the average parameters, are given in the Supplementary Information. Here, if we condition on *η_S_* = 0, then estimate of (*δ, α*) is

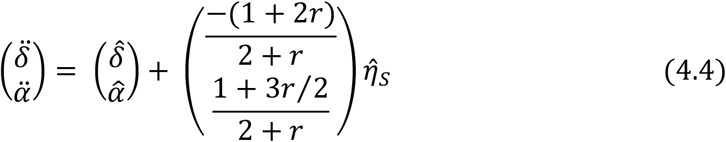

with variance

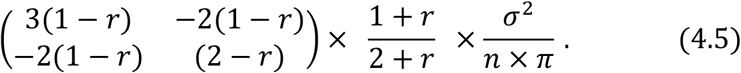

Note that 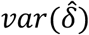 is an increasing function of *r*, but 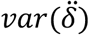 is a decreasing function of *r*. Thus, for estimating *δ*, one positively correlated double-proband sib-pair is less informative than two single proband sib-pairs without assuming *η_S_* equal to zero, but the opposite if *η_S_* is assumed to be zero. By contrast, without imputations, by regressing *Y*_ on (*G — G_S_*) and (*G* + *G_S_*), the fitted coefficient for (*G — G_S_*), denoted by 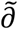, has expectation (*δ — η_S_*). When *G* is regressed on (*G — G_S_*) and (*G + G_S_*), the fitted coefficient of (*G* + *G_S_*), denoted by 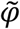, has expectation *δ* + *η_S_* + (4/3)*β*. It follows that 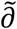 is an unbiased estimate of (*δ — η_S_*), and 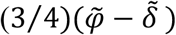 is an unbiased estimate of *β* + (3/2)*η_S_* = *α + η_S_*. The variance-covariance of 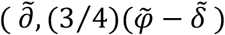 is approximately

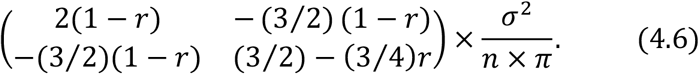

Thus, if *η_S_* ≠ 0, using 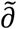 to estimate *δ* has a bias of —*η_S_*. By comparison, using 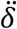 to estimate *δ*, the bias shrinks to — [(1 + 2*r*)/(2 + *r*)]*η_S_*. Even if *η_S_* is actually zero, 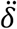 is more efficient, as measured by the inverse of the variance, than 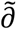. Assuming *η_S_* = 0, Fig. 4 shows the efficiency of estimating *δ*, as a function of *r*, using *n* double-proband sibpairs. Solid black line is without imputation of parental genotypes and solid red line is with imputations. The efficiency presented is relative to 2*n* single proband genotyped sibpairs without imputations. Notably, if *r* = 0, the efficiency of *n* double-proband sibpairs is statistically equivalent to *2n* single probands. Comparing the solid red line with the solid black line shows that, for *r* = (0, 0-1,0-2,0-3), 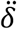 is (33,27,22,18)% more efficient than 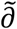.

**Figure 4:**
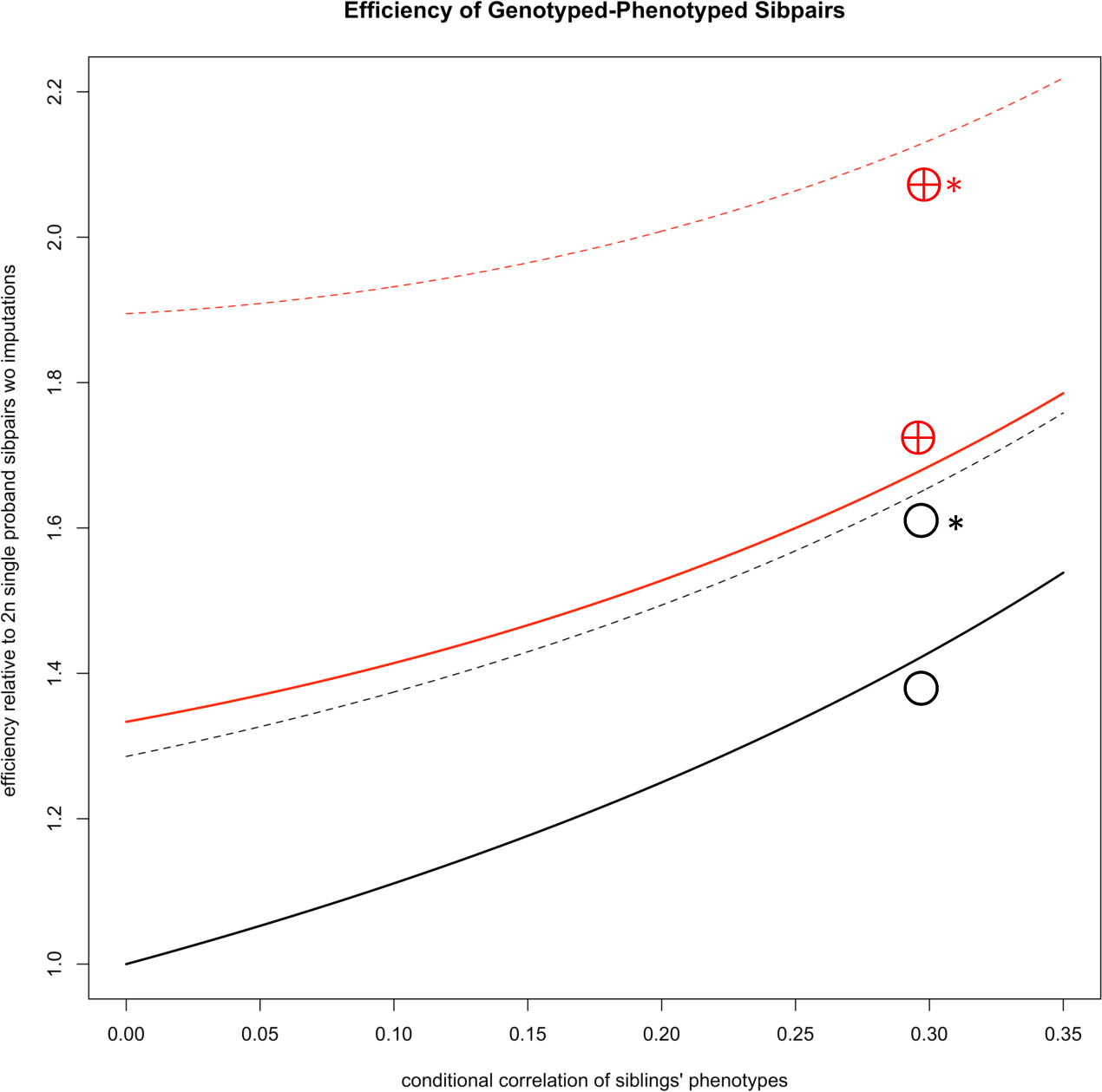
Efficiency of *n* double-proband genotyped sibpairs in estimating. ***δ***, under the assumption of *η_S_* = 0. The parental genotypes are assumed missing, *i.e.* the case in Fig 1e. Efficiency is displayed relative to 2*n* single-proband genotyped sibpairs and as a function of *r* = cor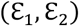. Black solid line (○) represents efficiency without imputation of parental genotypes. Red solid line (⊕) is efficiency with imputations. Black broken line (○*) is without imputation and augmented by 16n singletons. Red broken line (⊕*) is with imputations and augmented by 16n singletons.

While data from *singletons* (Fig. 1h), by themselves, cannot provide an unbiased estimate of *δ*, as an augmentation to the sibpair data, they can increase the efficiency of estimating *δ* and other parameters, a characteristic of multivariate meta-analysis. Without imputations, for *r* = (0,0-1,0-2,0-3), adding 16*n* singletons to *n* genotyped-phenotyped sibpairs increases the efficiency respectively from (1-0,1-11,1-25,1-43) to (1-29,1-37,1-49,1-66) (broken black line in Fig. 4), a percentage increase of (29,24,20,16). With imputations, efficiency increases from (1-33,1-41,1-53,1-68) to (1-89,1-93,2-01,2-13) (broken red line in Fig. 4), a percentage increase of (42, 37,31,27). As a consequence, when augmented by the singletons, for *r* = (0,0-1,0-2,0-3), imputing parental genotypes increases efficiency by a percentage of (47, 31, 34, 29) respectively (by comparing the broken red line with the broken black line). The 16-fold *singletons* versus double-proband sibpairs chosen for demonstration here is approximately the ratio seen in the UK Biobank samples. In Fig. 4, the broken black line is just below the solid red line. Indeed, if the sibpairs are augmented by an infinite number of singletons, then the two lines would coincide, *i.e.* the efficiency gain for estimating *δ* through IBD-based imputation of parental genotypes is the same as augmenting by practically an infinite number of singletons.

It is noted that our approach to sib-pairs can be extended naturally to incorporate sib-ships with three or more genotyped sibs^8^. In particular, with *k* genotyped sibs, on average 4(1 — 2^-*k*^) of the parental alleles can be deduced.

### Polygenic Scores and Assortative Mating

Consider a polygenic score of *T* SNPs:

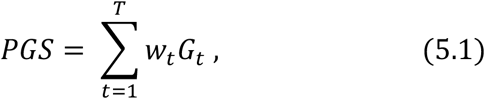

where *G_t_* and *w_t_* denote respectively genotype and weight of SNP *t*. When a person is not genotyped, the imputed polygenic score is

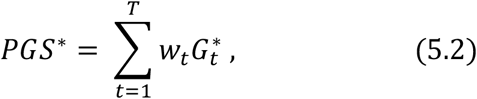

where each 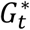 is imputed as before. Assuming the *T* SNPs are in linkage equilibrium with each other, then essentially all the previous results apply. For example, if the observed and imputed *PGSs* are jointly scaled so that *var(PGS)* = 1, then Table 1 gives the variance-covariance matrix of the observed and imputed *PGS* for various missing data patterns. Most importantly, if (3.7) holds for individual SNP genotypes, then it also holds for the polygenic scores since the variance-covariance matrix of the *PGSs* is just a weighted average of the variance-covariance matrixes of the individual SNP genotypes. If model (1.3) is generalized as

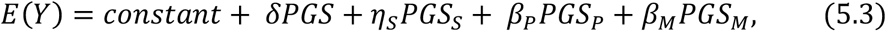

then estimate consistency will continue to apply with analysis performed with imputed polygenic scores. In practice, correlations, *i.e.* linkage disequilibrium (LD), between some of the SNPs are expected. However, as long as many SNPs contribute to the polygenic score and only a small fraction of the SNP pairs have non-negligible correlation, the effect on the imputations and estimates would be negligible. The phenomenon that requires consideration is assortative mating (ASM). For a trait with substantial assortative mating, contributing SNPs can become correlated regardless of their relative physical positions, an effect that is reduced, but usually not eliminated, by principal component (PC) adjustments^1^. Effects of ASM on genotype-phenotype associations are in general subtle and complicated, and have to be treated on a case by case basis. With imputations of parental genotypes based on genotypes of sib-pairs, ASM would lead to deviations between Σ_*X*_ and Σ_*col*_, *i.e.* violation of (3.7) and estimate consistency. However, as long as the trait is highly polygenic, *i.e.* the genetic component not dominated by a few variants, the estimates of *δ* and *η_S_* remain essentially unbiased. The estimate of *β* or *α* will be magnified by a multiplicative factor, but the degree of magnification is small unless the case is extreme. For example, if couples’ trait correlation is 0.30 and trait-PGS *R*^2^ is 0.35, the multiplicative factor is around 1.05. By contrast, if imputations are not performed and *Y* is regressed on *PGS* and *PGS_S_* only, the fitted coefficient of *PGS_S_* has expectation *η_S_* + (2/3)*β* under random mating. With ASM as described, the fitted coefficient has expectation around *η_S_* + −71 × *β* = *η_S_* + (1-065) × (2/3) × *β*. Thus IBD-based imputations actually reduce bias relative to no imputations. Most importantly, if deemed necessary, observed correlation of *PGS* and *PGS_S_*, which would go substantially above 0.5 with strong ASM, can be used to adjust the parameter estimates. It is noted that the bias referred to here is about using imputed data relative to having complete data. Even with complete data, as described previously^7^, as a consequence of ASM, estimates of *α* or *β* would capture, in addition to indirect effects, some confounding effects due to the parental *PGSs* being correlated with the part of the genetic component of the traits that is not captured by the *PGS* studied.

### Empirical Study

Using the developed methods, the effects of a EA polygenic score on EA, age-at-first-birth (AAFB)^10^, height (HT) and body-mass-index (BMI) are examined. The weights of 510,290 SNPs underlying this polygenic score are calculated (Supplementary Information) based on a GWAS meta-analysis that includes 608,402 samples, a subset of a larger set^2^, and includes 350,000 samples from UKB. A set of more than 39,000 probands from UKB with at least one sibling/parent genotyped is used to estimate the various effects of the polygenic score. The 350,000 individuals do not include any of the 39,000 probands or any third degree or higher relatives of the probands. Phenotypes are standardized for males and females separately to have variance one after adjusting for year-of-birth and 40 principal components. The number of probands for AAFB is about 40% of that for the other traits because AAFB information is only available for women from UKB and only applies to those who have children. This reduces sample size, but, in contrast to other sib-pair analyses that required both siblings to be phenotyped, our AAFB analysis includes 5216 probands, one-third of the total, with only genotyped male siblings.

After standardizing the polygenic scores so that those computed from observed genotypes have variance one, estimates of *δ,η_S_, β_M_*, and *β_P_* are obtained. After reparametrization, values of 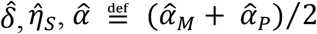, and 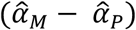 are displayed in Supplementary Table 2. For all four traits, 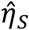 and 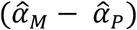 are not significantly different from zero. This does not mean *η_S_* is zero or that there are no parent-of-origin effects, only that they are not large enough to detect at our sample size. To simplify and to reduce variance, accepting the possibility of introducing some small bias, estimates 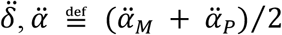, and 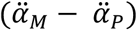 are computed conditioning on 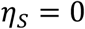 (Table 2). For all four traits, 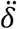 and 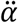 are highly significant (absolute effect size at least 5 times standard error (SE)). For EA, AAFB, BMI and HT, the estimated direct-population effect ratio, 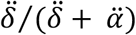, is 0.49, 0.66, 0.63 and 0.44 respectively. By comparison, for a different but related EA polygenic score applied to Icelandic data^7^, the corresponding ratio estimates are 0.70, 0.64, 0.72, and 0.42. The estimates are broadly consistent with the exception of EA. As *R*^2^, or variance explained is proportional to effect^2^, 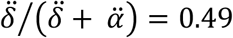 means the variance explained by the direct effect alone is only 0.49^2^ = 0.24 of the variance explained by the direct effect. In absolute terms, the population effect is estimated to explain (0.118 + 0.121)^2^ = 5.7% of the variance of EA, while it is only 0.118^2^ = 1.4% for the direct effect alone. Another striking result with the EA polygenic score is that its estimated direct effect for AAFB, 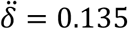, is higher than that for EA itself, 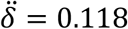. To make a more direct comparison, analysis for EA is recalculated using the same AAFB probands, and the estimated direct effect is 0.114. For these probands, correlation of EA and AAFB is 0.24, significant but not extremely high, indicating the polygenic score influences EA and AAFB mainly through separate causal paths, not purely affecting one trait through the other.

**Table 2.** Estimated effects of an EA polygenic scores. Estimates of direct effect 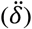, average parental effect 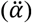, and the difference of the parental effects 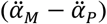, for the traits educational attainment (EA), age at first birth (AAFB), body mass index (BMI), and height (HT). Sibling effect *η_S_* is assumed to be zero. SE denotes standard error. Descriptions of the polygenic score and the UKB samples used are in the Supplementary Information.

| *trait* | $\ddot{\delta}$ | (SE) | $\ddot{\alpha}$ | (SE) | $\ddot{\alpha}_M - \ddot{\alpha}_P$ | (SE) |
| --- | --- | --- | --- | --- | --- | --- |
| EA | 0.118 | (0.008) | 0.121 | (0.007) | -0.017 | (0.026) |
| AAFB | 0.135 | (0.014) | 0.069 | (0.011) | 0.033 | (0.044) |
| BMI | -0.068 | (0.009) | -0.041 | (0.007) | 0.027 | (0.027) |
| HT | 0.04 | (0.007) | 0.052 | (0.007) | -0.017 | (0.027) |

It is noted that the singleton probands in the UKB data set cannot be used to augment the family analysis here because these singletons are part of the GWAS sample used to obtian the weights of the polygenic score.

## Discussion

We introduce Mendelian imputations as a tool to perform family-based association analysis. Conceptually, this is similar to multipoint linkage analysis performed with pedigrees that include deceased members. It is also related to familial imputations (also called in silico genealogy-based genotyping)^11,12^ and association by proxy^13,14^ where genotypes of relatives are used to associate with phenotypes of un-genotyped probands. Even though our general framework can also incorporate association by proxy, e.g. Fig. 1i to 1o, the main focus here is to disentangle various effects that contribute to the associations between a proband’s genotypes and phenotypes. As such, Mendelian imputations should also be applicable to family-based Mendelian randomization studies using genotyped sib-pairs^15^.

Mendelian imputations allow us to combine data with different missing data patterns in a single analysis, maximizing power. Moreover, even with one data type, Mendelian imputation increases flexibility and power. Specifically, for genotyped sib-pairs, it is shown that a genotyped sibling of the proband, even without phenotype, can be used. When both sibs are phenotyped, we can examine whether the genotypes have different effects on the sibs with respect to birth order or gender through direct or indirect effects. Non-linear imputations of parents not only increase power, but allow for the estimation of sibling nurturing effect, and enable unbiased estimation of the direct effect when the sibling nurturing effect is present. A recent manuscript^16^ proposed an imputation method that imputes each SNP without using IBD information inferred from neighbouring SNPs, *e.g.* when the two siblings are discordant homozygotes, then all four parental alleles can be inferred. While this method improves on not imputing at all, the gain is small relative to our method^8^, analogous to the difference between single-point and multipoint linkage analyses.

By applying the proposed method to UKB data, in addition to replicating observations previously reported based on Icelandic data, there are two thought provoking results. The low direct-associate effect ratio of the EA polygenic score on EA might be because many UK samples are included in the GWAS being used to construct the PGS, and thus part of the PGS’s predictive power with EA in other UKB samples could be population stratification effects that have not been eliminated by PC adjustments. It is known that the genetic components underling EA and AAFB have substantial overlap^17^, and the genetic component to EA was shown to have a stronger effect on AAFB of women than that of men in Iceland^18^. Despite that, the fact that the EA polygenic has a higher estimated direct effect on AAFB than EA is surprising. This highlights the complexity of the nature of the genetic variants influencing EA and reproductive traits. The issues raised here have important implications for how to interpret the population effects of a polygenic score, its portability and policy use. Family based analysis with more data and improved methodology could play a major role in the future research in this area.

## Acknowledgements

This work was supported by the Li Ka Shing Foundation. We thank the UK Biobank (application ID 11867). We thank Aysu Okbay for providing educational attainment summary statistics.

## References

1. Young, A. I., Benonisdottir, S., Przeworski, M. & Kong, A. Deconstructing the sources of genotype-phenotype associations in humans. Science. 365, 1396–1400 (2019).

2. Lee, J. J. et al. Gene discovery and polygenic prediction from a genome-wide association study of educational attainment in 1.1 million individuals. Nat. Genet. 50, 1112–1121 (2018).

3. Berg, J. J. et al. Reduced signal for polygenic adaptation of height in UK Biobank. Elife 8, 1–47 (2019).

4. Mostafavi, H. et al. Variable prediction accuracy of polygenic scores within an ancestry group. Elife 9, (2020).

5. Fulker, D. W., Cherny, S. S., Sham, P. C. & Hewitt, J. K. Combined linkage and association sib-pair analysis for quantitative traits. Am. J. Hum. Genet. 64, 259–267 (1999).

6. Jackson, D., Riley, R. & White, I. R. Multivariate meta-analysis: potential and promise. Stat. Med. 30, 2481–2498 (2011).

7. Kong, A. et al. The nature of nurture: Effects of parental genotypes. Science. 359, 424–428 (2018).

8. Young, A. I. et al. Mendelian imputation of parental genotypes for genome-wide estimation of direct and indirect genetic effects. BioRxiv (2020) doi:10.1101/185199.

9. Manichaikul, A. et al. Robust relationship inference in genome-wide association studies. Bioinformatics 26, 2867–2873 (2010).

10. Barban, N. et al. Genome-wide analysis identifies 12 loci influencing human reproductive behavior. Nat. Genet. 48, 1462–1472 (2016).

11. Kong, A. et al. Detection of sharing by descent, long-range phasing and haplotype imputation. Nat. Genet. 40, 1068–75 (2008).

12. Gudbjartsson, D. F. et al. A frameshift deletion in the sarcomere gene MYL4 causes early-onset familial atrial fibrillation. Eur. Heart J. ehw379 (2016).

13. Liu, J. Z., Erlich, Y. & Pickrell, J. K. Case--control association mapping by proxy using family history of disease. Nat. Genet. 49, 325 (2017).

14. Hujoel, M. L. A., Gazal, S., Loh, P.-R., Patterson, N. & Price, A. L. Liability threshold modeling of case--control status and family history of disease increases association power. Nat. Genet. 52, 541–547 (2020).

15. Brumpton, B. et al. Within-family studies for Mendelian randomization: avoiding dynastic, assortative mating, and population stratification biases. bioRxiv (2019) doi:10.1101/602516.

16. Hwang, L.-D. et al. Estimating indirect parental genetic effects on offspring phenotypes using virtual parental genotypes derived from sibling and half sibling pairs. BioRxiv (2020).

17. Mills, M. C., Tropf, F. C., Brazel, D. M., Zuydam, N. Van & Vaez, A. Identification of 370 loci for age at onset of sexual and reproductive behaviour, highlighting common aetiology with reproductive biology, externalizing behaviour and longevity. BioRxiv (2020).

18. Kong, A. et al. Selection against variants in the genome associated with educational attainment. Proc. Natl. Acad. Sci. U. S. A. 114, E727–E732 (2017).

