## Supplementary Information for "Family Analysis with Mendelian Imputations" (from another version of this preprint: version 1, 2020-07-03)

#### Linear Prediction

Let

$$Z = \begin{pmatrix} Z_1 \\ Z_2 \end{pmatrix} \qquad \text{(S.1)}$$

be a random vector with mean

$$\mu = \begin{pmatrix} \mu_1 \\ \mu_2 \end{pmatrix} \qquad \text{(S.2)}$$

and variance-covariance matrix

$$\Sigma = \begin{pmatrix} \Sigma_{11} & \Sigma_{12} \\ \Sigma_{21} & \Sigma_{22} \end{pmatrix}. \qquad \text{(S.3)}$$

The best linear (and unbiased) predictor of $Z_1$ given $Z_2$ is

$$\mu_1 + \Sigma_{12}\Sigma_{22}^{-1}(Z_2 - \mu_2). \qquad \text{(S.4)}$$

with variance

$$\Sigma_{11} - \Sigma_{12}\Sigma_{22}^{-1}\Sigma_{21}. \qquad \text{(S.5)}$$

For example, let $Z_1 = G_P$, and $Z_2 = (G, G_M)'$. Then $\mu_1 = (2p)$, $\mu_2 = (2p, 2p)'$, and

$$\Sigma_{12}\Sigma_{22}^{-1} = (\,1/2 \quad 0)\begin{pmatrix} 1 & 1/2 \\ 1/2 & 1 \end{pmatrix}^{-1} = (2/3 \quad -1/3\,).$$

It follows that the best linear predictor of $G_P$ given $G$ and $G_M$ is

$$2p + (2/3)G - (1/3)G_M - 2p(2/3) + 2p(1/3) = (2/3)G - (1/3)G_M + (4/3)p,$$

as given in (2.1) of the main text.

**Conditions for Estimate Consistency**

As noted in (3.7) of the main text,

$$\Sigma_X = \Sigma_{cov} \qquad \text{(S.6)}$$

is a key condition for *estimate consistency*. Through simulations and direct calculations, we have confirmed that the condition is satisfied with our proposed imputations. In particular, not only does Table 1 give the (scaled) variance-covariance matrix of the combination of actual and imputed genotypes under various missing-data patterns (scaled $\Sigma_X$), these same matrixes are also the scaled $\Sigma_{cov}$. Here we provide the mathematical principles underlying (S.6). For notation, follow (S.1) to (S.3) and let $Z_1$ be the unobserved genotypes, $Z_1^*$ be its imputed version, and $Z_2$ be the observed genotypes. Furthermore, for simplicity, we centre the $Z's$ so that $\mu$ is zero, e.g. $Z_M = G_M - 2p$, etc.

Equation (S.6) is satisfied IFF

$$cov(Z_2, Z_1^*) = cov(Z_2, Z_1) \qquad \text{(S.6a)}$$

and

$$var(Z_1^*) = cov(Z_1^*, Z_1). \qquad \text{(S.6b)}$$

First tackle (S.6a). Start with $Z_1^*$ being the linear prediction. From (S.4),

$$Z_1^{*L} = \Sigma_{12}\Sigma_{22}^{-1}Z_2$$

and

$$cov(Z_2, Z_1^{*L}) = cov(Z_2, \Sigma_{12}\Sigma_{22}^{-1}Z_2) = var(Z_2)(\Sigma_{12}\Sigma_{22}^{-1})' = \Sigma_{22}\Sigma_{22}^{-1}\Sigma_{21} = \Sigma_{21}.$$

Thus (S.6a) is satisfied by linear imputation. The imputations we propose utilize the observed genotypes plus IBD information deduced from the IBS statuses of neighbouring markers plus phasing. Other non-linear imputations utilize less information. For example, for genotyped sibpairs where both parents are not genotyped, one can use deduced IBD number (0, 1, 2) between sibs, but do not use phasing information (again deduced from neighbouring markers) to determine which is the sharing allele in the case of double-heterozygotes[1]. The method by Hwang et al[2] does not use information from neighbouring markers at all. Even then there is still some non-linear information to be gained, even though much reduced. For example, if the siblings' genotypes are discordant homozygotes, then we know all four parental alleles are observed, and if one sibling is homozygote and the other is heterozygote, then we know at least 3 parental alleles are observed. These different non-linear imputations can be considered as conditional expectations of $Z_1$ given $Z_2$ plus different degrees of auxiliary IBD information. As such,

$$cov(Z_2, Z_1^*) = E\left[Z_2 Z_1^{*'}\right] = E[Z_2 E(Z'_1|Z_2)] = E[Z_2 Z_1'] = cov(Z_2, Z_1) = cov(Z_2, Z_1^{*L}).$$

In other words, the covariances between the imputed and observed genotypes are the same for linear imputations and various form of non-linear conditional expectation imputations. This can be seen by comparing Table 1 with Supplementary Table 1. But the correlations are not the same because $var(Z_1^*)$ are different for the different imputations. In particular, higher values of $var(Z_1^*)$ correspond to more information introduced by the imputations (compare Table 1 and Supplementary Table 1 for cases where non-linear imputations are possible) and lower correlations between the imputed and observed genotypes. Expressing $Z_1^*$ as conditional expectation, we have

$$cov(Z_1^*, Z_1) = E[E(Z_1|Z_2)Z_1'] = E\{E[E(Z_1|Z_2)Z_1'|Z_2]\} = E\{E[E(Z_1|Z_2)E(Z_1'|Z_2)|Z_2]\}$$
$$= E\{E(Z_1|Z_2)E(Z_1'|Z_2)\} = var\big(E(Z_1|Z_2)\big) = var(Z_1^*),$$

giving (S.6b).

**Double-proband sib-pair analysis**

Double-proband refers to the case when both genotyped sibs are also phenotyped. Starting by not assuming symmetry, we can fit (4.1a) and (4.2a) separately to obtain estimates of sib specific parameters. However, these parameter estimates are correlated because $\mathcal{E}_1$ and $\mathcal{E}_2$ are in general correlated. To obtain standard errors for estimates that average over the sibs and/or conditioning on certain parameters equal to zero, the joint variance-covariance of all the parameter estimates is needed. In an empirical study, bootstrapping the sib-pairs is one solution. An alternative, which also applies to theoretical investigations, is reparameterization of the $Y$'s. With $Y_- \overset{\text{def}}{=} Y_1 - Y_2$ and $Y_+ \overset{\text{def}}{=} Y_1 + Y_2$, we have

$$Y_- = constant + \vec{G}'\left[\vec{\theta}_1 - \vec{\vartheta}\right] + \mathcal{E}_-, \qquad \text{(S.7a)}$$

and

$$Y_+ = constant + \vec{G}'\left[\vec{\theta}_1 + \vec{\vartheta}\right] + \mathcal{E}_+, \qquad \text{(S.7b)}$$

where $\mathcal{E}_- = \mathcal{E}_1 - \mathcal{E}_2$ and $\mathcal{E}_+ = \mathcal{E}_1 + \mathcal{E}_2$. Let $cor(\mathcal{E}_1, \mathcal{E}_2) = r$. As long as $\vec{G}$ only accounts for a small fraction of the variances of the $Y$'s, $cor(\mathcal{E}_-, \mathcal{E}_+)$ is either exactly zero, when $Y_1$ and $Y_2$ are symmetric, or approximately zero. Furthermore, assuming $Y_1$ and $Y_2$ are scaled to have the same variance,

$$var(\mathcal{E}_-) \cong (2-2r)\sigma^2,\ var(\mathcal{E}_+) \cong (2+2r)\sigma^2, \quad \text{(S.8)}$$

where $\sigma^2$ is the average of $var(\mathcal{E}_1)$ and $var(\mathcal{E}_2)$, and the difference between $var(\mathcal{E}_1)$ and $var(\mathcal{E}_2)$ is assumed to be zero or negligibly small. The mathematics and methods established in the main text for the single proband case can be applied to $Y_-$ and $Y_+$ separately obtaining uncorrelated estimates (exactly or approximately) of two parameter vectors. They can then be combined to provide estimates of up to eight parameters. Here we examine the case where only the siblings are genotyped, one of the most common data-types. Because neither parent is genotyped, by imputing the parents jointly, our method regresses $Y_-$ and $Y_+$ on $\vec{X} = (G, G_S, G^*_{PM})$. Here

$$\Sigma_X = \begin{pmatrix} 1 & 1/2 & 1 \\ 1/2 & 1 & 1 \\ 1 & 1 & 3/2 \end{pmatrix}\pi \quad \text{and } \Sigma_X^{-1} = \begin{pmatrix} 4 & 2 & -4 \\ 2 & 4 & -4 \\ -4 & -4 & 6 \end{pmatrix}/\pi. \quad \text{(S.9)}$$

where $\pi = 2p(1-p)$. Fitted coefficients from regressing $Y_-$ and $Y_+$ on $\vec{X}$ , denoted by $\hat{\tau}$ and $\hat{\upsilon}$ have expectations

$$\tau \stackrel{\text{def}}{=} \begin{pmatrix} \delta_1 \\ \eta_{S1} \\ \beta_1 \end{pmatrix} - \begin{pmatrix} \eta_{S2} \\ \delta_2 \\ \beta_2 \end{pmatrix} = \begin{pmatrix} \delta_1 - \eta_{S2} \\ \eta_{S1} - \delta_2 \\ \beta_1 - \beta_2 \end{pmatrix} \text{and } \upsilon \stackrel{\text{def}}{=} \begin{pmatrix} \delta_1 \\ \eta_{S1} \\ \beta_1 \end{pmatrix} + \begin{pmatrix} \eta_{S2} \\ \delta_2 \\ \beta_2 \end{pmatrix} = \begin{pmatrix} \delta_1 + \eta_{S2} \\ \eta_{S1} + \delta_2 \\ \beta_1 + \beta_2 \end{pmatrix}. \quad \text{(S.10)}$$

respectively. The respective variances are

$$\frac{var(\mathcal{E}_-)}{n}\Sigma_X^{-1} \cong \frac{2(1-r)\sigma^2}{n}\Sigma_X^{-1}. \quad \text{(S.11)}$$

and

$$\frac{var(\mathcal{E}_+)}{n}\Sigma_X^{-1} \cong \frac{2(1+r)\sigma^2}{n}\Sigma_X^{-1}. \quad \text{(S.12)}$$

Through linear transformation,

$$\begin{pmatrix} \delta \\ \eta_S \\ \alpha \\ \delta_- \\ \eta_{S-} \\ \alpha_- \end{pmatrix} \stackrel{\text{def}}{=} \begin{pmatrix} \phi \\ \\ \psi \end{pmatrix} = A \begin{pmatrix} \tau \\ \\ \upsilon \end{pmatrix} \quad \text{(S.13)}$$

where

$$A = \begin{pmatrix} 1/4 & -1/4 & 0 & 1/4 & 1/4 & 0 \\ -1/4 & 1/4 & 0 & 1/4 & 1/4 & 0 \\ -1/8 & 1/8 & 0 & 1/8 & 1/8 & 1/2 \\ 1/2 & 1/2 & 0 & 1/2 & -1/2 & 0 \\ 1/2 & 1/2 & 0 & -1/2 & 1/2 & 0 \\ 1/4 & 1/4 & 1 & -1/4 & 1/4 & 0 \end{pmatrix} \stackrel{\text{def}}{=} \begin{pmatrix} A_{11} & A_{12} \\ A_{21} & A_{22} \end{pmatrix}. \quad \text{(S.14)}$$

Note that $\phi$ and $\psi$ are vectors with length 3, and $A_{11,}$ $A_{12}, A_{21}, A_{22}$ are $3 \times 3$ matrices. Estimate of $\phi$ is

$$\hat{\phi} = \begin{pmatrix} \hat{\delta} \\ \hat{\eta}_S \\ \hat{\alpha} \end{pmatrix} = A_{11}\,\hat{\tau} + A_{12}\,\hat{\upsilon}. \qquad \text{(S.15)}$$

Based on the fact that $\hat{\tau}$ and $\hat{\upsilon}$ are uncorrelated with variances given in (S.11) and (S.12), the variance-covariance matrix of $\hat{\phi}$ can be shown to be

$$\cong \begin{pmatrix} 2+r & 1+2r & -\left(\frac{3}{2}+r\right) \\ 1+2r & 2+r & -\left(1+\frac{3}{2}r\right) \\ -\left(\frac{3}{2}+r\right) & -\left(1+\frac{3}{2}r\right) & \frac{3}{2}+\frac{5}{4}r \end{pmatrix} \times \frac{\sigma^2}{n\times\pi}. \qquad \text{(S.16)}$$

If we condition on $\eta_S = 0$, then estimate of $(\delta, \alpha)$ is

$$\begin{pmatrix} \ddot{\delta} \\ \ddot{\alpha} \end{pmatrix} = \begin{pmatrix} \hat{\delta} \\ \hat{\alpha} \end{pmatrix} + \begin{pmatrix} \frac{-(1+2r)}{2+r} \\ \frac{1+3r/2}{2+r} \end{pmatrix} \hat{\eta}_S \qquad \text{(S.17)}$$

with variance

$$\begin{pmatrix} 3(1-r) & -2(1-r) \\ -2(1-r) & (2-r) \end{pmatrix} \times \frac{1+r}{2+r} \times \frac{\sigma^2}{n\times\pi}. \qquad \text{(S.18)}$$

By contrast, without imputations, by regressing $Y_-$ on $(G - G_S)$ and $(G + G_S)$, the fitted coefficient for $(G - G_S)$, denoted by $\tilde{\partial}$, has expectation $(\delta - \eta_S)$. When $Y_+$ is regressed on on $(G - G_S)$ and $(G + G_S)$, the fitted coefficient of $(G + G_S)$, denoted by $\tilde{\varphi}$, has expectation $\delta + \eta_S + (4/3)\beta$. It follows that $\tilde{\partial}$ is an unbiased estimate of $(\delta - \eta_S)$, and $(3/4)(\tilde{\varphi} - \tilde{\delta}\,)$is an unbiased estimate of $\beta + (3/2)\eta_S = \alpha + \eta_S$. The variance-covariance of $(\,\tilde{\partial}, (3/4)(\tilde{\varphi} - \tilde{\delta}\,)$ is

$$\begin{pmatrix} 2(1-r) & -(3/2)\,(1-r) \\ -(3/2)(1-r) & (3/2)-(3/4)r \end{pmatrix} \times \frac{\sigma^2}{n\times\pi}. \qquad \text{(S.19)}$$

Similarly, the variances of the estimates of the difference parameters, which are uncorrelated with $\hat{\phi}$, can be shown to be

$$var\begin{pmatrix} \hat{\delta}_1 - \hat{\delta}_2 \\ \hat{\eta}_{S1} - \hat{\eta}_{S2} \\ \hat{\alpha}_1 - \hat{\alpha}_2 \end{pmatrix} \cong \begin{pmatrix} 8-4r & 4-8r & -6+4r \\ 4+8r & 8-4r & -4+6r \\ -6+4r & -4+6r & 6-5r \end{pmatrix} \times \frac{\sigma^2}{n\times\pi}. \qquad \text{(S.20)}$$

### Empirical Study

*Polygenic Score.* Educational attainment polygenic scores were computed using meta-analysis results from the study of Lee et al after excluding UKBB individuals with at least one 1st degree relative in the data and 23andMe samples. As our study of direct and indirect effects is based on siblings and parent-offspring pairs, UKBB individuals with 1st degree relatives in the data were exluded from the meta-analysis to avoid confounding and/or overfitting. Number of individuals used in the meta-analysis varies with SNP, ranging from 300,026 individuals to 606,402 individuals (mean = 556,264; SD = 64.6). The polygenic scores were computed from direct genotype calls of 510,290 high-quality biallelic SNPs with MAF>1% and non-ambigous nucleotides. SNP effects were adjusted for LD using LDpred[2] version 1.08. LD structure was approximated using a subset of 10,090 UKBB samples of WB descent with no close relative in UKBB (>3rd degree for all possible pairs). We choose fraction of causal variants with respect to prediction accuracy in a subset of 7,472 UKBB samples of WB descent that have at least one sibling in the data but are not closely related to each other (>3rd degree for all possible pairs). EA polygenic scores were computed for siblings and parent-offspring pairs in UKBB by summing over weighted alleles of the 510,290 SNPs using the LD adjusted effects as weights[3]. This EA polygenic score explains 5.97% of phenotypic variance in the set of 7,472 individuals with at least one sibling.

*Samples Analysed.* We extracted the genotypes for that subsample of the UK Biobank, removing SNPs with missingness above 5%. We used KING[20] with the '--related --degree 1' options to infer the sibling and parent-offspring relations within that set of individuals. We identified 157 duplicates/monozygotic twins and removed one from each pair from further analyses. We identified 17,296 families with at least two siblings, giving a total of 19,329 sibling pairs. The maximum number of siblings in a family was 6, and 913 families had more than two siblings. We used age and sex information to determine the father/mother in each inferred parent-offspring relation, requiring parents to have a reported age at least 12 years higher than their inferred child; parent-offspring relations with a lower reported age difference were removed from further analyses. We identified 4,418 families with at least one parent and one child genotyped; 736 families had at least one child and the father genotyped but not the mother genotyped; 2,798 families had at least one child and the mother but not the father genotyped; 893 families had at least one child and both parents genotyped. We identified 31 families with two children and both parents genotyped, ‘

1. Young, A. I. *et al.* Mendelian imputation of parental genotypes for genome-wide estimation of direct and indirect genetic effects. *BioRxiv* (2020) doi:10.1101/185199.

2. Hwang, L.-D. *et al.* Estimating indirect parental genetic effects on offspring phenotypes using virtual parental genotypes derived from sibling and half sibling pairs. *BioRxiv* (2020).
3. Vilhjálmsson, B. J. *et al.* Modeling Linkage Disequilibrium Increases Accuracy of Polygenic Risk Scores. *Am. J. Hum. Genet.* **97**, 576–592 (2015).

a)

$$\begin{array}{c} \begin{matrix} \boldsymbol{G} & \boldsymbol{G_S} & \boldsymbol{G_P} & \boldsymbol{G_M} \end{matrix} \\ \begin{pmatrix} 1 & 1/2 & 1/2 & 1/2 \\ 1/2 & 1 & 1/2 & 1/2 \\ 1/2 & 1/2 & 1 & 0 \\ 1/2 & 1/2 & 0 & 1 \end{pmatrix} \end{array}$$

b)

$$\begin{array}{c} \begin{matrix} \boldsymbol{G} & \boldsymbol{G_S^*} & \boldsymbol{G_M} & \boldsymbol{G_P} \end{matrix} \\ \begin{pmatrix} 1 & 1/2 & 1/2 & 1/2 \\ 1/2 & 1/2 & 1/2 & 1/2 \\ 1/2 & 1/2 & 1 & 0 \\ 1/2 & 1/2 & 0 & 1 \end{pmatrix} \end{array}$$

c)

$$\begin{array}{c} \begin{matrix} \boldsymbol{G} & \boldsymbol{G_S^*} & \boldsymbol{G_P} & \boldsymbol{G_M^*} \end{matrix} \\ \begin{pmatrix} 1 & 1/2 & 1/2 & 1/2 \\ 1/2 & 1/3 & 1/2 & 1/6 \\ 1/2 & 1/2 & 1 & 0 \\ 1/2 & 1/6 & 0 & 1/3 \end{pmatrix} \end{array}$$

d)

$$\begin{array}{c} \begin{matrix} \boldsymbol{G} & \boldsymbol{G_S^*} & \boldsymbol{G_P^*} & \boldsymbol{G_M} \end{matrix} \\ \begin{pmatrix} 1 & 1/2 & 1/2 & 1/2 \\ 1/2 & 1/3 & 1/6 & 1/2 \\ 1/2 & 1/6 & 1/3 & 0 \\ 1/2 & 1/2 & 0 & 1 \end{pmatrix} \end{array}$$

e)

$$\begin{array}{c} \begin{matrix} \boldsymbol{G} & \boldsymbol{G_S} & \boldsymbol{G_P^*} & \boldsymbol{G_M^*} \end{matrix} \\ \begin{pmatrix} 1 & 1/2 & 1/2 & 1/2 \\ 1/2 & 1 & 1/2 & 1/2 \\ 1/2 & 1/2 & 1/3 & 1/3 \\ 1/2 & 1/2 & 1/3 & 1/3 \end{pmatrix} \end{array}$$

f)

$$\begin{array}{c} \begin{matrix} \boldsymbol{G} & \boldsymbol{G_S} & \boldsymbol{G_P} & \boldsymbol{G_M^*} \end{matrix} \\ \begin{pmatrix} 1 & 1/2 & 1/2 & 1/2 \\ 1/2 & 1 & 1/2 & 1/2 \\ 1/2 & 1/2 & 1 & 0 \\ 1/2 & 1/2 & 0 & 1/2 \end{pmatrix} \end{array}$$

g)

$$\begin{array}{c} \begin{matrix} \boldsymbol{G} & \boldsymbol{G_S} & \boldsymbol{G_P^*} & \boldsymbol{G_M} \end{matrix} \\ \begin{pmatrix} 1 & 1/2 & 1/2 & 1/2 \\ 1/2 & 1 & 1/2 & 1/2 \\ 1/2 & 1/2 & 1/2 & 0 \\ 1/2 & 1/2 & 0 & 1 \end{pmatrix} \end{array}$$

h)

$$\begin{array}{c} \begin{matrix} \boldsymbol{G} & \boldsymbol{G_S^*} & \boldsymbol{G_P^*} & \boldsymbol{G_M^*} \end{matrix} \\ \begin{pmatrix} 1 & 1/2 & 1/2 & 1/2 \\ 1/2 & 1/4 & 1/4 & 1/4 \\ 1/2 & 1/4 & 1/4 & 1/4 \\ 1/2 & 1/4 & 1/4 & 1/4 \end{pmatrix} \end{array}$$

i)

$$\begin{array}{c} \begin{matrix} \boldsymbol{G^*} & \boldsymbol{G_S} & \boldsymbol{G_P} & \boldsymbol{G_M} \end{matrix} \\ \begin{pmatrix} 1/2 & 1/2 & 1/2 & 1/2 \\ 1/2 & 1 & 1/2 & 1/2 \\ 1/2 & 1/2 & 1 & 0 \\ 1/2 & 1/2 & 0 & 1 \end{pmatrix} \end{array}$$

j)

$$\begin{array}{c} \begin{matrix} \boldsymbol{G^*} & \boldsymbol{G_S} & \boldsymbol{G_P} & \boldsymbol{G_M^*} \end{matrix} \\ \begin{pmatrix} 1/3 & 1/2 & 1/2 & 1/6 \\ 1/2 & 1 & 1/2 & 1/2 \\ 1/2 & 1/2 & 1 & 0 \\ 1/6 & 1/2 & 0 & 1/3 \end{pmatrix} \end{array}$$

k)

$$\begin{array}{c} \begin{matrix} \boldsymbol{G^*} & \boldsymbol{G_S} & \boldsymbol{G_P^*} & \boldsymbol{G_M} \end{matrix} \\ \begin{pmatrix} 1/3 & 1/2 & 1/6 & 1/2 \\ 1/2 & 1 & 1/2 & 1/2 \\ 1/6 & 1/2 & 1/3 & 0 \\ 1/2 & 1/2 & 0 & 1 \end{pmatrix} \end{array}$$

l)

$$\begin{array}{c} \begin{matrix} \boldsymbol{G^*} & \boldsymbol{G_S} & \boldsymbol{G_P^*} & \boldsymbol{G_M^*} \end{matrix} \\ \begin{pmatrix} 1/4 & 1/2 & 1/4 & 1/4 \\ 1/2 & 1 & 1/2 & 1/2 \\ 1/4 & 1/2 & 1/4 & 1/4 \\ 1/4 & 1/2 & 1/4 & 1/4 \end{pmatrix} \end{array}$$

m)

$$\begin{array}{c} \begin{matrix} \boldsymbol{G^*} & \boldsymbol{G_S^*} & \boldsymbol{G_P} & \boldsymbol{G_M} \end{matrix} \\ \begin{pmatrix} 1/2 & 1/2 & 1/2 & 1/2 \\ 1/2 & 1/2 & 1/2 & 1/2 \\ 1/2 & 1/2 & 1 & 0 \\ 1/2 & 1/2 & 0 & 1 \end{pmatrix} \end{array}$$

n)

$$\begin{array}{c} \begin{matrix} \boldsymbol{G^*} & \boldsymbol{G_S^*} & \boldsymbol{G_P} & \boldsymbol{G_M^*} \end{matrix} \\ \begin{pmatrix} 1/4 & 1/4 & 1/2 & 0 \\ 1/4 & 1/4 & 1/2 & 0 \\ 1/2 & 1/2 & 1 & 0 \\ 0 & 0 & 0 & 0 \end{pmatrix} \end{array}$$

o)

$$\begin{array}{c} \begin{matrix} \boldsymbol{G^*} & \boldsymbol{G_S^*} & \boldsymbol{G_P^*} & \boldsymbol{G_M} \end{matrix} \\ \begin{pmatrix} 1/4 & 1/4 & 0 & 1/2 \\ 1/4 & 1/4 & 0 & 1/2 \\ 0 & 0 & 0 & 0 \\ 1/2 & 1/2 & 0 & 1 \end{pmatrix} \end{array}$$

**Supplementary Table 1: Linear imputation and resulting variance-covariance matrices**. Each matrix shows the variance-covariance structure within the nuclear family given observed and linearly imputed unobserved genotypes. $G$ symbols the probands genotype, $G_s$ symbols the sibling’s genotype, $G_P$ symbols the fathers genotype and $G_M$ symbols the mothers genotype. * denotes an unobserved genotype that is linearly imputed from observed genotypes.

| *trait* | $\hat{\delta}$ | *(SE)* | $\hat{\eta}_s$ | *(SE)* | $\hat{\alpha}$ | *(SE)* | $\hat{\alpha}_M - \hat{\alpha}_P$ | *(SE)* |
|---|---|---|---|---|---|---|---|---|
| EA | 0.117 | (0.012) | -0.001 | (0.013) | 0.126 | (0.011) | -0.023 | (0.102) |
| AAFB | 0.139 | (0.019) | 0.019 | (0.019) | 0.064 | (0.017) | 0.144 | (0.149) |
| BMI | -0.068 | (0.013) | 0.002 | (0.013) | -0.042 | (0.011) | 0.078 | (0.105) |
| HT | 0.026 | (0.013) | -0.018 | (0.014) | 0.063 | (0.012) | 0.116 | (0.116) |

**Supplementary Table 2. Estimated effects of an EA polygenic scores.** Estimates of direct effect, sibling effect, average parental effect, and the difference of the parental effects, for the traits educational attainment (EA), age at first birth (AAFB), body mass index (BMI), and height (HT). SE denotes standard error.
